# Receptor activity modifying protein modulation of parathyroid hormone-1 receptor function and signalling

**DOI:** 10.1101/2024.04.16.588673

**Authors:** Paris Avgoustou, Ameera B. A. Jailani, Aditya J. Desai, Dave J. Roberts, Ewan R. Lilley, Grant W. Stothard, Timothy M. Skerry, Gareth O. Richards

## Abstract

Receptor activity-modifying proteins (RAMPs) are known to modulate the pharmacology and function of several G protein-coupled receptors (GPCRs), including the parathyroid hormone 1 receptor (PTH1R). However, the precise effects of different RAMPs on PTH1R signalling and trafficking remain poorly understood. Here we investigated the impact of RAMP2 and RAMP3 on PTH1R function using a range of PTHand PTH-related protein (PTHrP)-derived ligands.

FRET imaging revealed that PTH1R preferentially interacts with RAMP2 and, to a lesser extent, RAMP3, but not RAMP1. Interestingly, RAMP3 co-expression resulted in reduced cell surface expression of PTH1R, suggesting a potential role in receptor trafficking or internalization. The presence of RAMP2 significantly enhanced PTH1R-mediated cAMP accumulation, β-arrestin recruitment, and calcium signalling in response to PTH (1–34), PTHrP (1-34), PTH (1-84), and the PTH (1-17) analogue ZP2307. In contrast, RAMP3 co-expression attenuated or completely abolished those responses.

We found that full-length PTHrP analogues, PTHrP (1-108) and PTHrP (1-141), exhibited lower potency and efficacy than PTHrP (1-34) in activating PTH1R. RAMP2 significantly increased potency and/or efficacy when compared to PTH1R alone cells, while RAMP3 significantly reduced these responses. Antibody-capture scintillation proximity assays demonstrated that RAMP2 differentially modulates G protein activation by PTH1R in a ligand-dependent manner, with PTH (1-34) and PTHrP (1-34) inducing distinct patterns of G protein subtype activation.

These findings highlight the complex role of RAMPs in regulating PTH1R signalling and trafficking, revealing differential effects of RAMP2 and RAMP3 on receptor function. The data suggest that targeting the PTH1R/RAMP2 complex may be a promising strategy for developing novel bone anabolic therapies by leveraging biased agonism and functional selectivity. Further research using physiologically relevant models is needed to elucidate the therapeutic potential of this approach.

## Introduction

The G-protein coupled receptors (GPCRs), a superfamily of seven transmembrane domaincontaining receptors that consists of more than 800 members, are the largest family of membrane-bound proteins. GPCRs are involved in many physiological and pathophysiological processes (1) and are among the most numerous targets for drug development (∼35% of all FDA approved drugs, ∼700 drugs) (1, 2).

The parathyroid hormone receptor-1 (PTH1R) is a member of this family and is more specifically a class B GPCR, with a crucial role in a wide range of physiological systems including calcium homeostasis (in normal life, pregnancy, and lactation), skeletal development, and bone turnover. Pathological actions of the PTH1R include involvement in the pathophysiology of osteoporosis, hypoparathyroidism (3, 4), humoral hypercalcaemia of malignancy and increasing numbers of malignancies (5). These effects occur as a result of the PTH1R’s ability to bind and transduce signals mediated by two cognate ligands. Parathyroid hormone (PTH) and PTH-related peptide (PTHrP) are the two endogenous agonists that lead to the activation of PTH1R and mediate its pleiotropic functions (4). Although the PTH1R was classically thought to signal through the Gα_s_/cAMP pathway (activation) (4), it can also activate a number of other second-messenger cascades including Gα_q_/calcium influx (6), Gα_i_/cAMP pathway (inhibition) (7) and β-arrestins (8). Further complexity of this ligand/receptor system was added when PTH1R was found to interact with members of the receptor-activity-modifying protein (RAMP) family (9–11). Mammalian RAMPs (RAMP1, RAMP2, and RAMP3) are single transmembrane domain proteins known to interact with several GPCR family members altering and regulating receptor function and pharmacology (12–14) but with no ligand binding activities alone. RAMPs regulate ligand selectivity for a small number of GPCRs (Figure 1a) (15) and intracellular trafficking of rather more receptors putatively (Figure 1b) (16). Moreover, it was shown that RAMPs are involved in the regulation of GPCRs internalization and recycling (Figure 1c) (17, 18). In addition, it has been shown that RAMPs can modulate receptor downstream signalling when responding to the same ligands (19) (Figure 1d).

**Figure 1:**
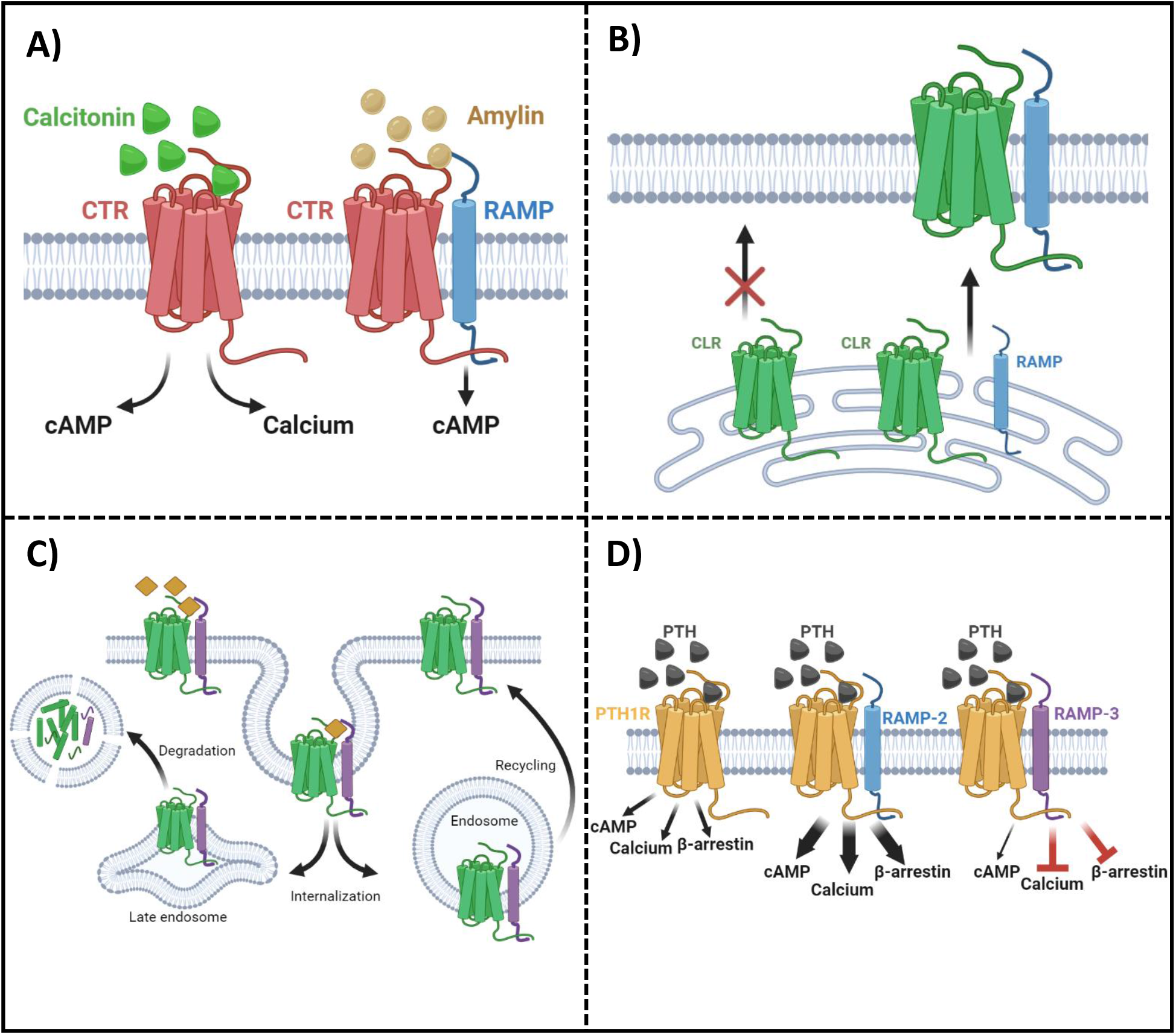
Summary of RAMP functions. a) Ligand selectivity: RAMPs were originally discovered in connection with their ability to alter the ligand selectivity of the calcitonin-like receptor and calcitonin receptor. Here expression of a RAMPs changes the phenotype of a calcitonin receptor so that instead of binding with calcitonin it binds with amylin, transducing different intracellular signalling pathways. **b) Receptor trafficking:** RAMPs are essential for trafficking of some receptors, which cannot localise at the cell surface alone. Calcitonin receptor like receptor (CLR) therefore only functions when trafficked with one of the three RAMPs. **c) Internalization and recycling:** With other receptors trafficking is possible without RAMP association, but it is suggested that there are alterations in the rate and kinetics of transport. Also, recent studies have shown that RAMPs alter the recycling rate of GPCRs. RAMP3 was responsible for retaining the AM2 receptor (RAMP3/CLR) intracellularly after agonist stimulation and for the rapid recycling of the atypical chemokine receptor ACKR3. **d)**

Most of the studies to date have focused on the interaction between PTH1R and RAMP2 (9, 10), although some studies include data on interactions of PTH1R with RAMP3 (11, 20). Nemec et al. in 2022, has shown that RAMP2 alters the PTH1R signalling in agonist dependent manner, with most significant increase in the PTH-mediated Gα_i3_ signalling sensitivity. Furthermore, RAMP2 caused an increase in both PTH (1–34)and PTHrP(1–34)-triggered βarrestin recruitment to the PTH1R (11).

The physiological consequences of PTH1R/RAMP interaction are still unclear. Evidence from RAMP2 knockout mice (RAMP2+/-) showed a decrease in PTH1R expression as well as a dampened response on serum phosphate concentration after systemic parathyroid hormone (PTH) administration, although it is worth mentioning that in this study a very large dose of PTH (500μg/kg) was used (21). In the same study, placental dysfunction and defects in arterial remodelling were observed in RAMP2+/mice that was not associated with RAMP2/CLR receptor complex, suggesting a possible physiological role of PTH1R/RAMP2 receptor complex (21).

In this study we have confirmed and added to the observations by Nemec at al. and expand on the repertoire of PTH derived ligands and the signalling consequences of both PTH1R/RAMP2 and PTH1R/RAMP3 interactions (Figure 1d).

### Intracellular consequences/signalling

RAMPs alter the intracellular consequences of the same ligand binding to the same receptor. Here we show that RAMP2 causes significant increases in second messenger activation (cAMP, calcium and β-arrestin) when compared to PTH1R alone, while RAMP3 causes significant reduction or complete inhibition of such processes.

## Results

### PTH1R and RAMPs interactions and cell surface trafficking

To identify which RAMPs, have potential for functional interactions with the PTH1R, we used a sensitized Fluorescent Resonance Energy Transfer (FRET) technique to determine receptor/RAMP interactions closer than 10nm at the cell surface in COS-7 cells transfected with different RAMPs and receptors. We also used FRET based stoichiometric analysis (22) to determine the fraction of receptor and RAMPs in the FRET complex.

FRET was quantified at the cell surface using membrane ROIs as shown in figure 2a. This showed that PTH1R interacts with RAMP2 and to a lesser extent with RAMP3, however no interaction was observed with RAMP1 (Figure 2a, 2b). The relative stoichiometry between PTH1R/RAMP2 and RAMP3 also differed (Figure 2c), suggesting that PTH1R and RAMP2 formed ∼1:1 complex, however, PTH1R and RAMP3 formed ∼1:2 complex (Figure 2c).

**Figure 2:**
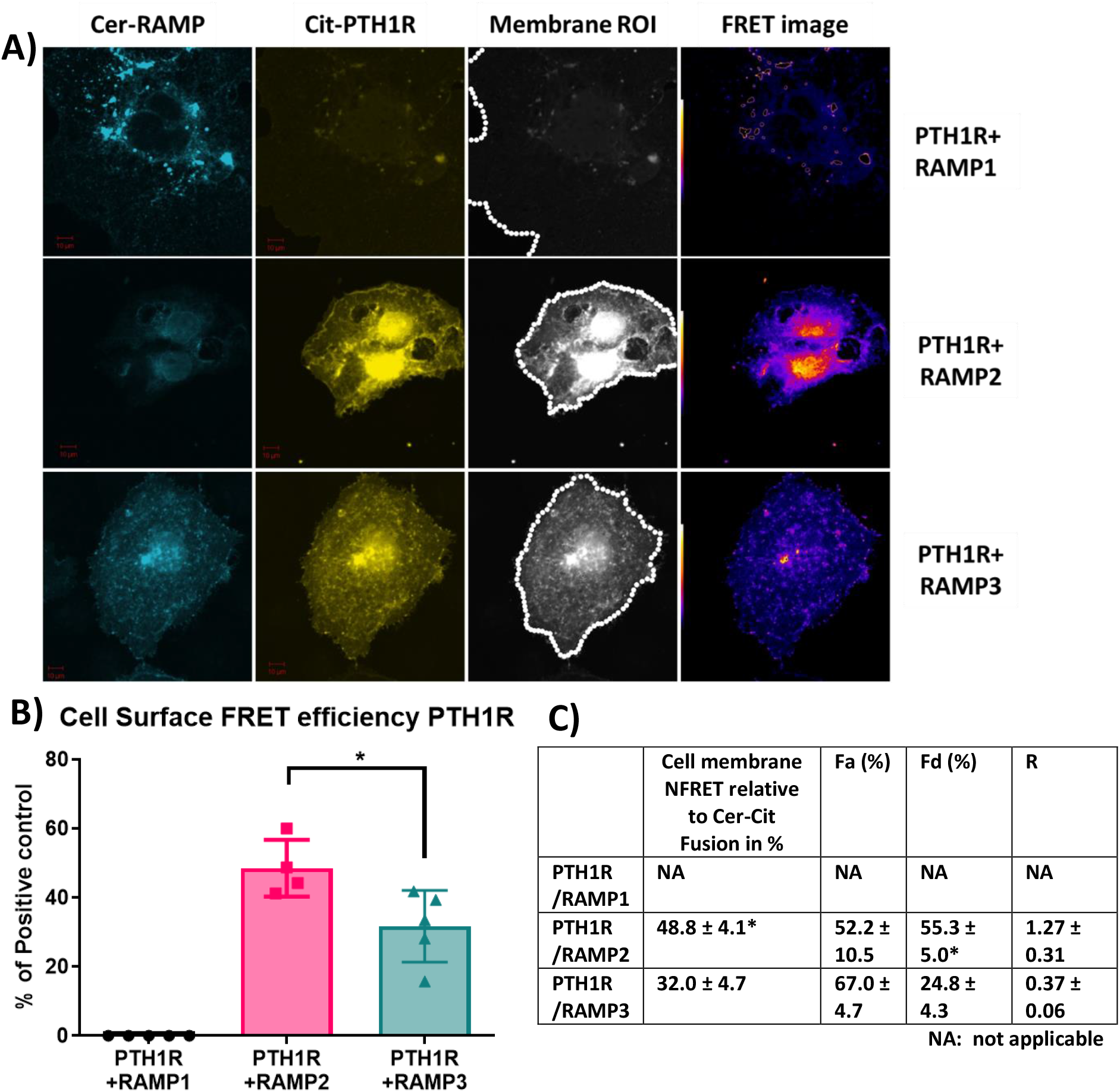
Membrane localisation of PTH1R and RAMPs. FRET imaging of COS-7 cells transfected with Cerulean-RAMPs 1-3 and Citrine-PTH1R combinations. Analysis of cell surface FRET was performed using a series of 50-pixel regions of interest (ROI) constructed around the Citrine-receptor raw image to cover the entire cell membrane. Images are representative of 4-6 replicate measurements. **a)** Confocal images of RAMP1-3 (Cerulean) PTH1R (Citrine) in COS-7 cell. **b)** FRET efficiencies of RAMP1-3 with PTH1R represented as % of the maximum FRET calculated using the Cerulean-Citrine fusion construct. **c)** Mean values for NFRET and the FRET stoichiometric constants for various RAMP and receptor combinations on the cell surface. Fa= fraction of GPCR (acceptor) in FRET complex. Fd= fraction of RAMP (donor) in FRET complex. R= molar ratio of acceptor to donor. NA indicates no detectable FRET. The data represent mean ± SEM, n=4. * p< 0.05 and ** p<0.001, MannWhitney test.

To support if RAMP interactions alter cell surface trafficking of PTH1R, we performed cell-based ELISA for the PTH1R, in the presence and absence of RAMPs (23). These studies were performed in stably expressing cells, as described in methods section. We observed that the presence of RAMP1 and RAMP2 did not alter the cell surface translocation of the PTH1R, compared to PTH1R cells. However, when co-expressed with RAMP3, cell surface levels of PTH1R were comparable to the parental cell line (CHO-K1) suggesting that RAMP3 may play a role in intracellular retention of PTH1R (Figure 3).

**Figure 3:**
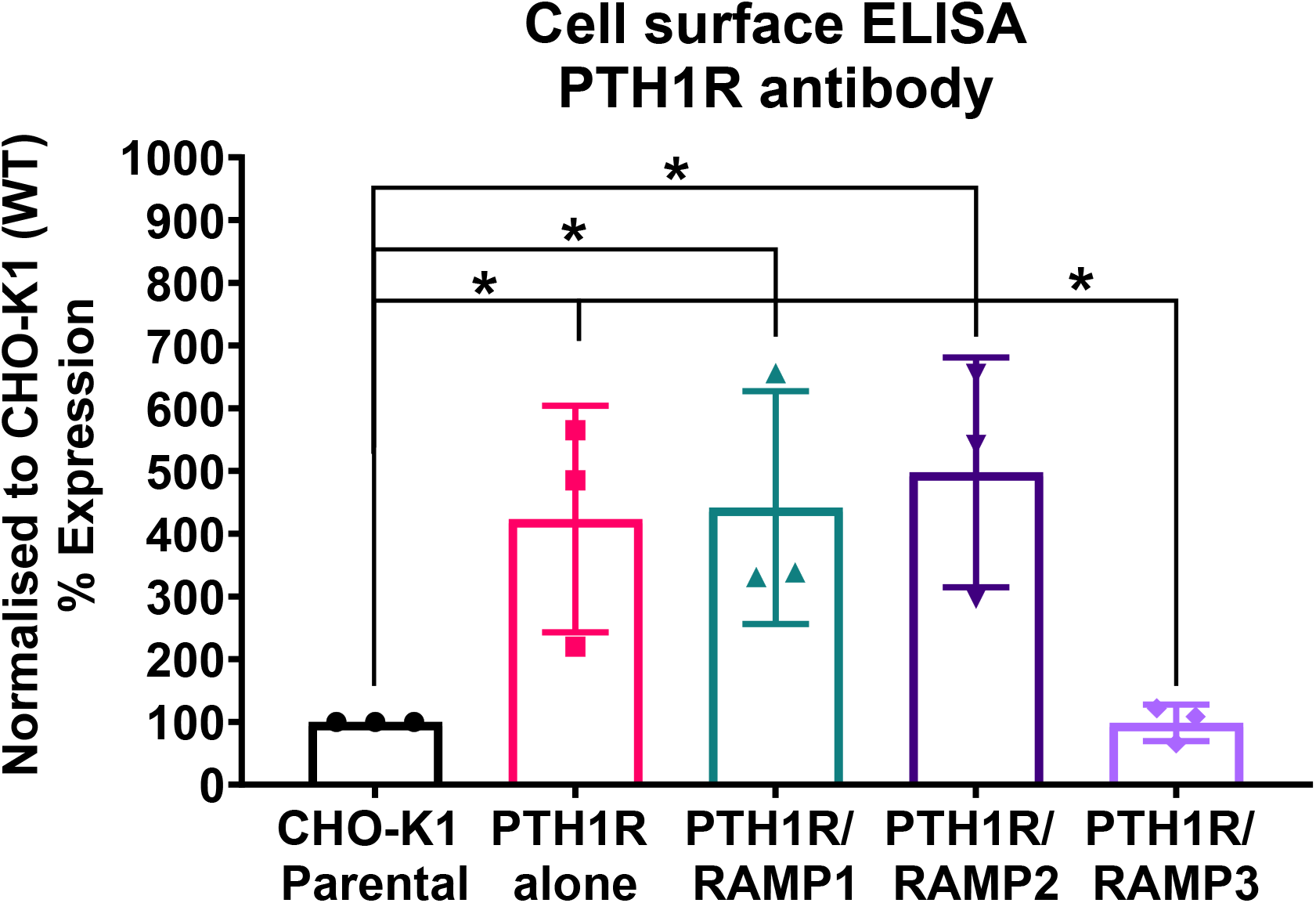
Cell-based ELISA against PTH1R, in the presence and absence of RAMPs. Cell surface ELISA was performed on cells stably expressing PTH1R alone, PTH1R/RAMP1, PTH1R/RAMP2, PTH1R/RAMP3 and the parental cell line (CHO-K1), against PTH1R antibody. Expression was normalised to CHO-K1 parental cell line. Data are derived from 3 replicate measurements in 3 independently replicated studies. Data are presented as mean ± SD. Comparisons were analysed using unpaired Student’s t-test, * p< 0.05.

### Consequences of RAMP2 interaction on PTH1R G-protein response to ligand activation

To explore the consequences of PTH1R/RAMP2 interaction, we used antibody-capture scintillation proximity assays (SPA) to measure the spectra of activation of individual Gproteins (Gα_i_, Gα_q_ and Gα_s_) by the PTH1R alone or in combination with RAMP2 (Figures 2, 3). These studies were controlled for levels of receptor expression (using radioligand binding studies and western blot, data not shown) and confirmed by cell surface expression ELISA disclosure (Figure 3).

In COS-7 cells membrane preparations expressing PTH1R alone, we assessed the effects of varying concentrations of PTH (1–34) and PTHrP (1–34) in the absence of RAMPs. PTH (1–34) induced a greater maximal activation (efficacy) of Gα_s_ (39%) and Gα_i_ (67%) compared to PTHrP(1–34) and a detectable activation in Gα_q_ that is absent with PTHrP (all p<0.05) (Figure 4a, 4c & 4e).

**Figure 4:**
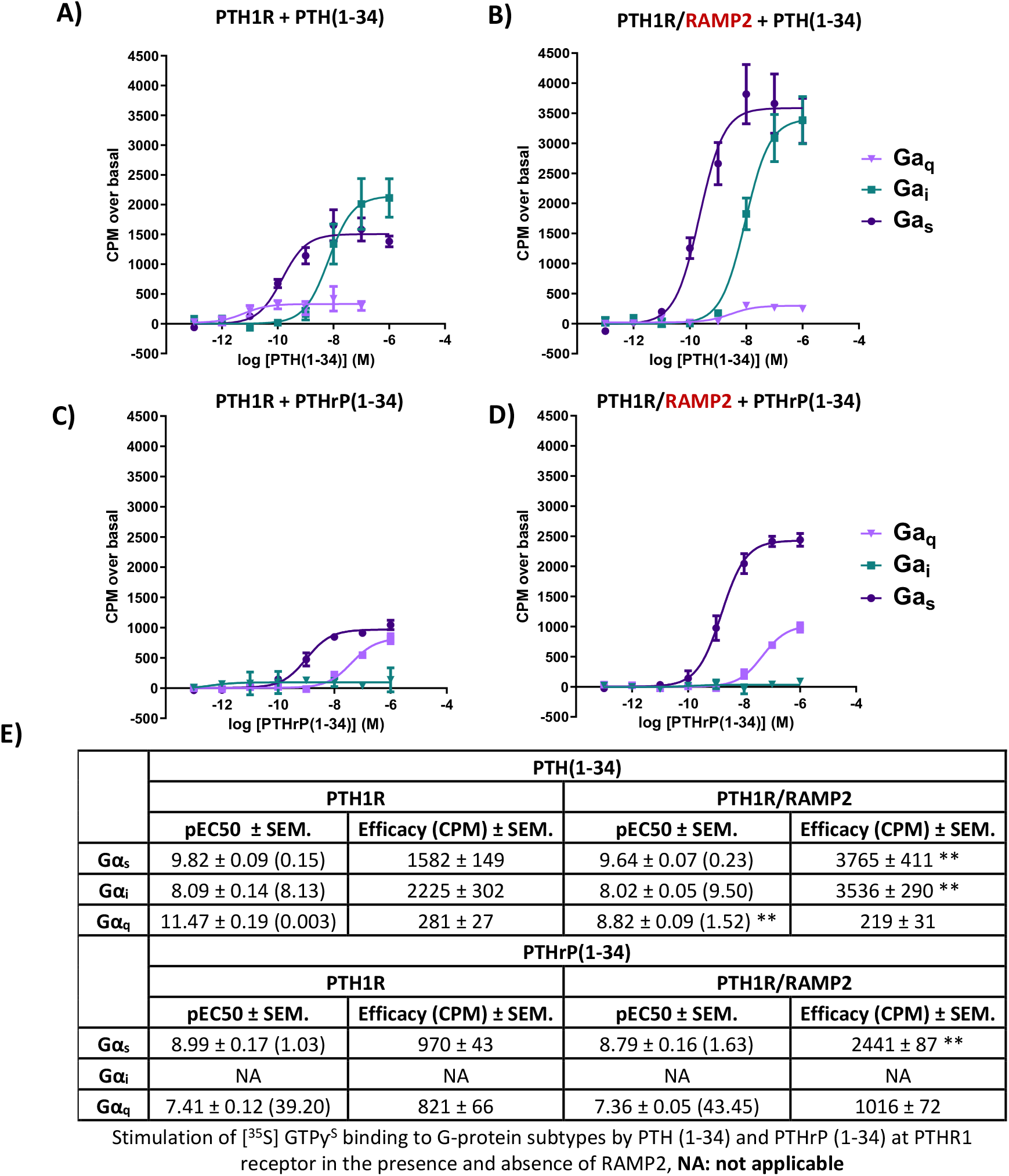
Antibody-capture scintillation proximity assays (SPA). Dose response curves constructed from experiments in which different concentrations of ligand (PTH (1–34) or PTHrP (1–34)) were incubated with 10µg of membrane preparations from COS-7 cells transfected either with PTH1R alone or PTH1R and RAMP2. Disclosure of G-protein activation was performed by addition of [^35^S]GTPɣ^S^ and scintillant beads to allow measurement. **a)** Responses of membranes from cells transfected with PTH1R to PTH (1–34). **b)** Responses of membranes from cells transfected with PTH1R/RAMP2 to PTH (1–34). **c)** Responses of the same membranes as a, transfected with PTH1R, to PTHrP (1–34). **d)** Responses of the same membranes as b, transfected PTH1R/RAMP2, to PTHrP (1–34). **e)** Potency (pEC50) and efficacy of G-protein activation derived from the curves obtained in SPA studies. Data are derived from curves constructed from 3 independently replicated studies, each consisting of 2 replicate measurements at each of 8 ligand concentrations. Table data shown as -log values ± SEM with nM values in brackets. Comparisons were analysed using ANOVA with Bonferroni post-hoc analysis (*P<0.05, **P<0.01 ***P<0.001).

We then assessed the effects of RAMP2 on PTH1R G-protein response to PTH (1–34) and PTHrP(1–34) activation. The interaction of PTH1R with RAMP2 increased PTH (1–34)stimulated maximal activation (efficacy) of Gα_s_ (by 140%) and Gα_i_ (by 60%), without changing Gα_q_ (Figure 4a, 4b & 4e) compared with the same ligand acting on the receptor alone. There were no changes in potency (EC_50_) for Gα_s_ and Gα_i_ activation, but PTH-stimulated Gα_q_ potency was significantly reduced in the PTH1R associated with RAMP2 compared with PTH1R alone. In contrast, PTHrP(1–34) induced a different pattern of changes in the same membrane preparations expressing PTH1R and RAMP2 compared to PTH1R alone. PTHrP-stimulated Gα_s_ efficacy was increased by ∼150% without changes in Gα_i_ or Gα_q_ or any changes in potency (Figure 4c, 4d & 4e).

### RAMP2 modulates receptor functionality in a ligand-dependent manner

Following on from the interaction and G protein activation studies we decided to investigate the effects of RAMP2 and RAMP3 on PTH1R second messenger activation in response to PTH (1–84), PTH (1–34), PTHrP (1–34), PTHrP (1–108) and PTHrP (1–141) and cyclic PTH (1–17), to expand on the repertoire of PTH derived ligands (Figure 5).

**Figure 5:**
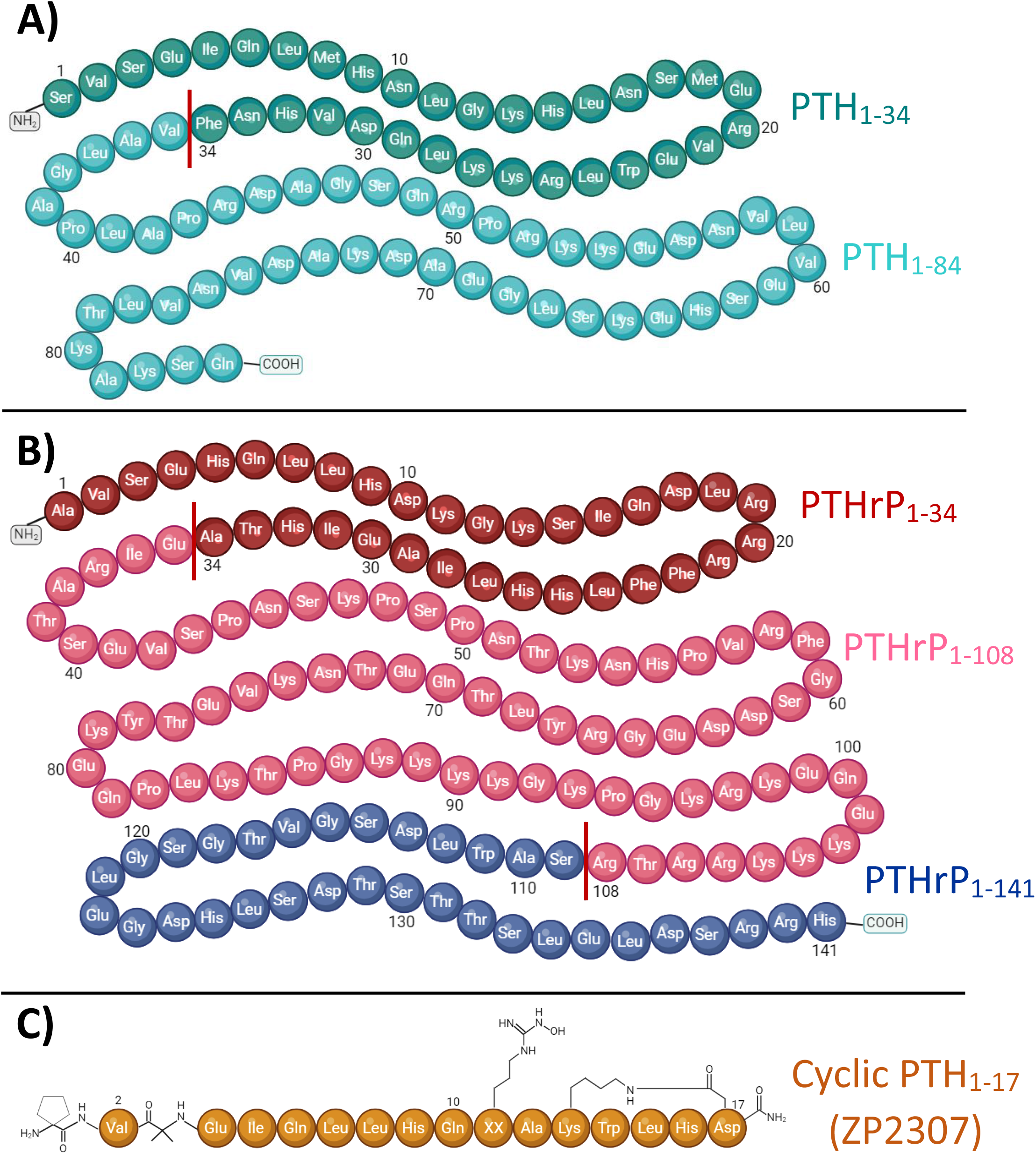
Primary sequences of PTH and PTHrP analogues. **a)** Amino acid sequence of human PTH_1-34_ and PTH_1-84_. The line separates the PTH_1-34_ amino acid segment from the full-length peptide (PTH_1-84_). **b)** Amino acid sequence of human PTHrP_1-34_ and PTHrP_1-108_ and PTHrP_1-141_. The lines separate the PTHrP_1-34_ and PTHrP_1-108_ amino acid segments from the full-length peptide PTHrP_1-141_. **c)** Amino acid sequence of PTH novel human cyclic PTH_1-17_ analogue (ZP2307).

### PTH (1–34)

PTH1R/RAMP2 cells showed a significant increase in both potency and maximal response (efficacy) to PTH (1–34) mediated cAMP accumulation, compared to PTH1R alone (Figure 6). In contrast, PTH1R/RAMP3 cells showed a significant reduction in potency but no changes in efficacy compared to PTH1R alone (Figure 6). Even though there was no significant difference in the potency of PTH (1–34) to recruit β-arrestin between PTH1R alone and PTH1R/RAMP2 cells there a was a significant increase in the efficacy of the ligand (PTH (1–34)) in the RAMP2transfected cells (Figure 6). RAMP3-transfected cells had no detectable response to PTH (1-34) in the β-arrestin assays. Similarly, when the same cells were used to measure calcium influx changes, a significant difference in efficacy was shown between PTH1R alone and PTH1R/RAMP2 cells but no significant change in potency (Figure 6). RAMP3-transfected cells had no response to PTH (1-34) in all the assays we performed. To assess the effects of RAMP2 in Gα_i_ response, the cells were treated with PTX (pertussis toxin) prior performing cAMP accumulation assays. In these assays the RAMP2 transfected cells showed a significant increase in both the potency and efficacy of PTH (1-34), compared to PTH1R alone cells. RAMP3 had no significant effect (Figure 6).

**Figure 6:**
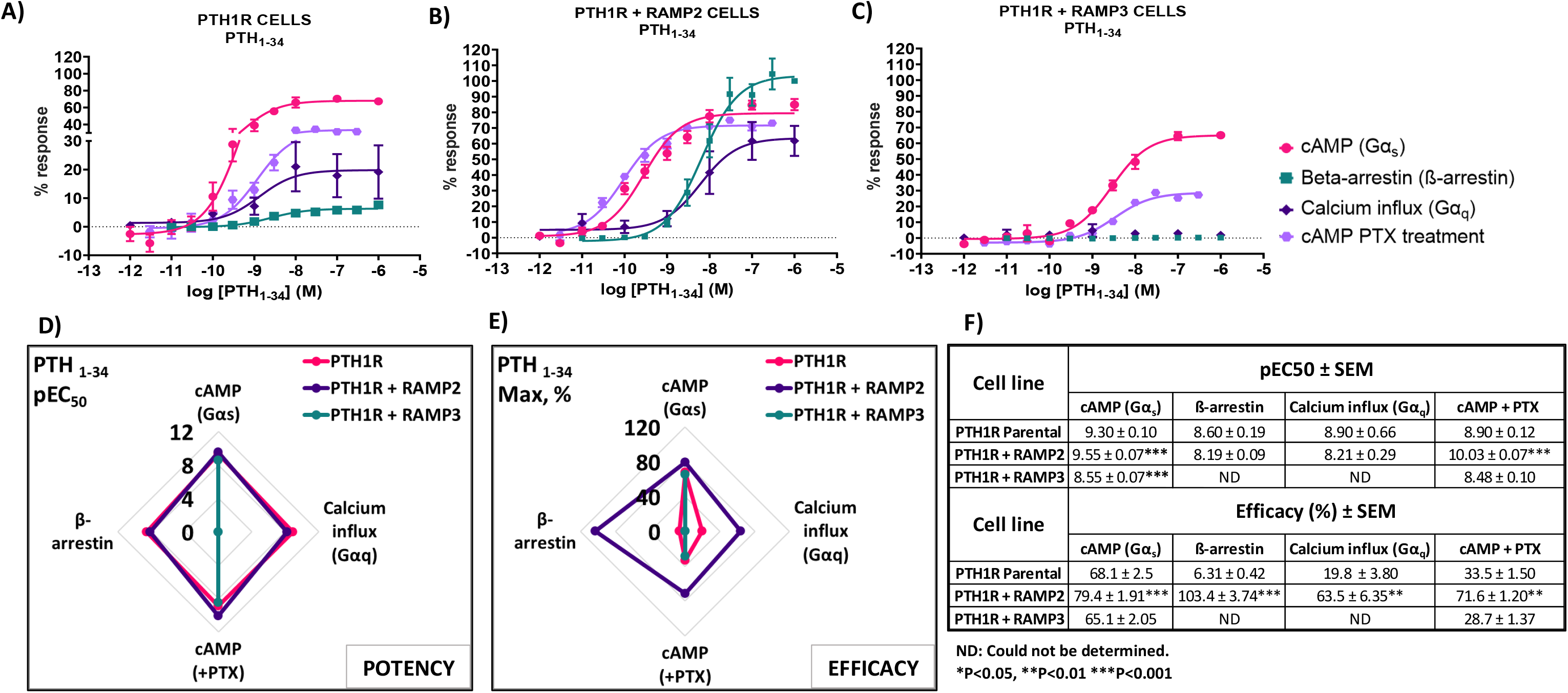
Consequences of RAMP2 and RAMP3 in the potency and efficacy of PTH (1-34) in different functional assays. Dose response curves of PTH (1-34) in different second messenger pathways, in cells overexpressing **a)** PTH1R alone, **b)** PTH1R with RAMP2 and **c)** PTH1R with RAMP3. Spider diagrams of the **d)** the potency and **e)** the efficacy of PTH (1-34) in those assays as extracted from the dose response curves. **f)** Table summarising the potency and the efficacy of PTH (1-34) and the statistical differences between the cells. Data are derived from curves constructed from at least 3-4 independently replicated studies, each consisting of 2 replicate measurements at each of 11 ligand concentrations. Data were analysed using comparison of fits (Graph Pad Prism) for non-linear regression curves, three-parameter logistic curve (*P<0.05, **P<0.01 ***P<0.001). All curves were expressed as a % of the positive controls/maximal response. Controls: cAMP studies forskolin (100μM), calcium influx studies ATP (100μM), β-arrestin-2 recruitment studies maximal response at highest dose (1μM).

### PTHrP (1-34)

When PTH1R RAMP2 cells were stimulated with PTHrP (1-34) a significant increase in potency but not efficacy was shown in cAMP accumulation, compared to PTH1R alone (Figure 7). On the other hand, transfection with RAMP3 showed a significant reduction in potency but no changes in efficacy compared to PTH1R alone (Figure 7). Similarly with the mediated effects of PTH (1-34) on β-arrestin, a significant increase in the efficacy of the ligand (PTHrP (1-34)) in the RAMP2 transfected cells was shown (Figure 7). In addition, RAMP3 transfected cells had no response to PTHrP (1-34) in the β-arrestin assays (Figure 7). When stimulated with PTHrP (1-34), only cells transfected with RAMP2 showed an increase in calcium influx, while PTH1R alone and PTH1R RAMP3 cells showed no response (Figure 7). In Gα_i_ response, RAMP2 caused a significant increase in both potency and efficacy of PTHrP (1-34), compared to PTH1R alone. On the other hand, RAMP3 caused a significant reduction in in both potency and efficacy (Figure 7).

**Figure 7:**
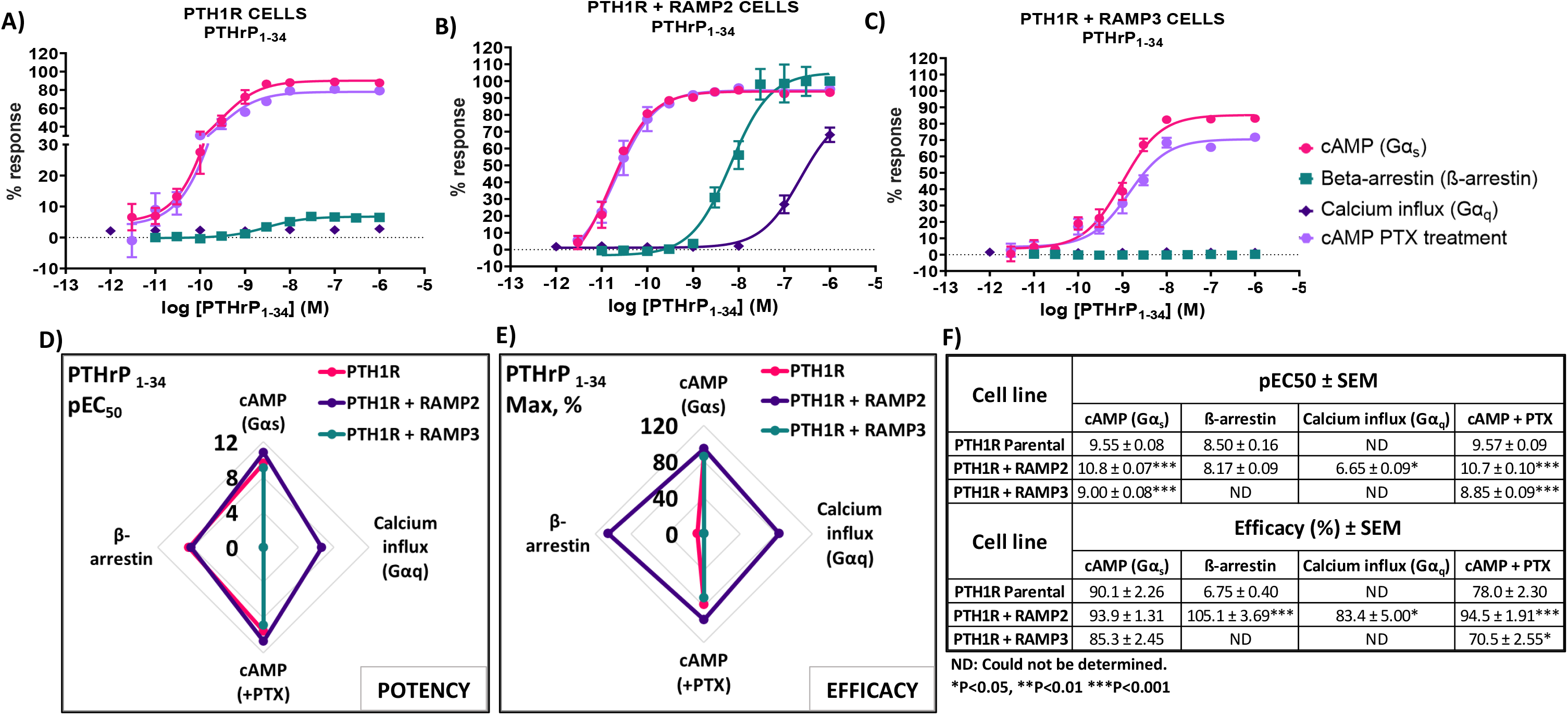
Consequences of RAMP2 and RAMP3 in the potency and efficacy of PTHrP (1-34) in different functional assays. Dose response curves of PTHrP (1-34) in different second messenger pathways, in cells overexpressing **a)** PTH1R alone, **b)** PTH1R with RAMP2 and **c)** PTH1R with RAMP3. Spider diagrams of the **d)** the potency and **e)** the efficacy of PTHrP (1-34) in those assays as extracted from the dose response curves. **f)** Table summarising the potency and the efficacy of PTHrP (1-34) and the statistical differences between the cells. Data are derived from curves constructed from at least 3-4 independently replicated studies, each consisting of 2 replicate measurements at each of 11 ligand concentrations. Data were analysed using comparison of fits (Graph Pad Prism) for non-linear regression curves, three-parameter logistic curve (*P<0.05, **P<0.01 ***P<0.001). All curves were expressed as a % of the positive controls/maximal response. Controls: cAMP studies forskolin (100μM), calcium influx studies ATP (100μM), β-arrestin-2 recruitment studies maximal response at highest dose (1μM).

### PTH (1-84)

While using the intact biologically active 84 amino acid peptide (PTH (1-84)), similar effects where shown. RAMP2 transfected cells (PTH1R RAMP2) showed a significant increase in potency in cAMP accumulation (Gα_s_) and in Gα_i_ responses, but no changes in efficacy (Figure 8). On the other hand, a significant increase in efficacy of PTH (1-84) was shown in β-arrestin. But no changes in potency (Figure 8). Similarly, with PTHrP (1-34), only cells transfected with RAMP2 showed an increase in calcium influx, while PTH1R alone and PTH1R RAMP3 cells showed no response when stimulated with PTH (1-84) (Figure 8). Transfection with RAMP3 resulted in a significant decrease in both potency and efficacy PTH (1-84) in in cAMP accumulation (Gα_s_), in efficacy in Ga_i_ and no response in β-arrestin when compared with PTH1R alone cells (Figure 8).

**Figure 8:**
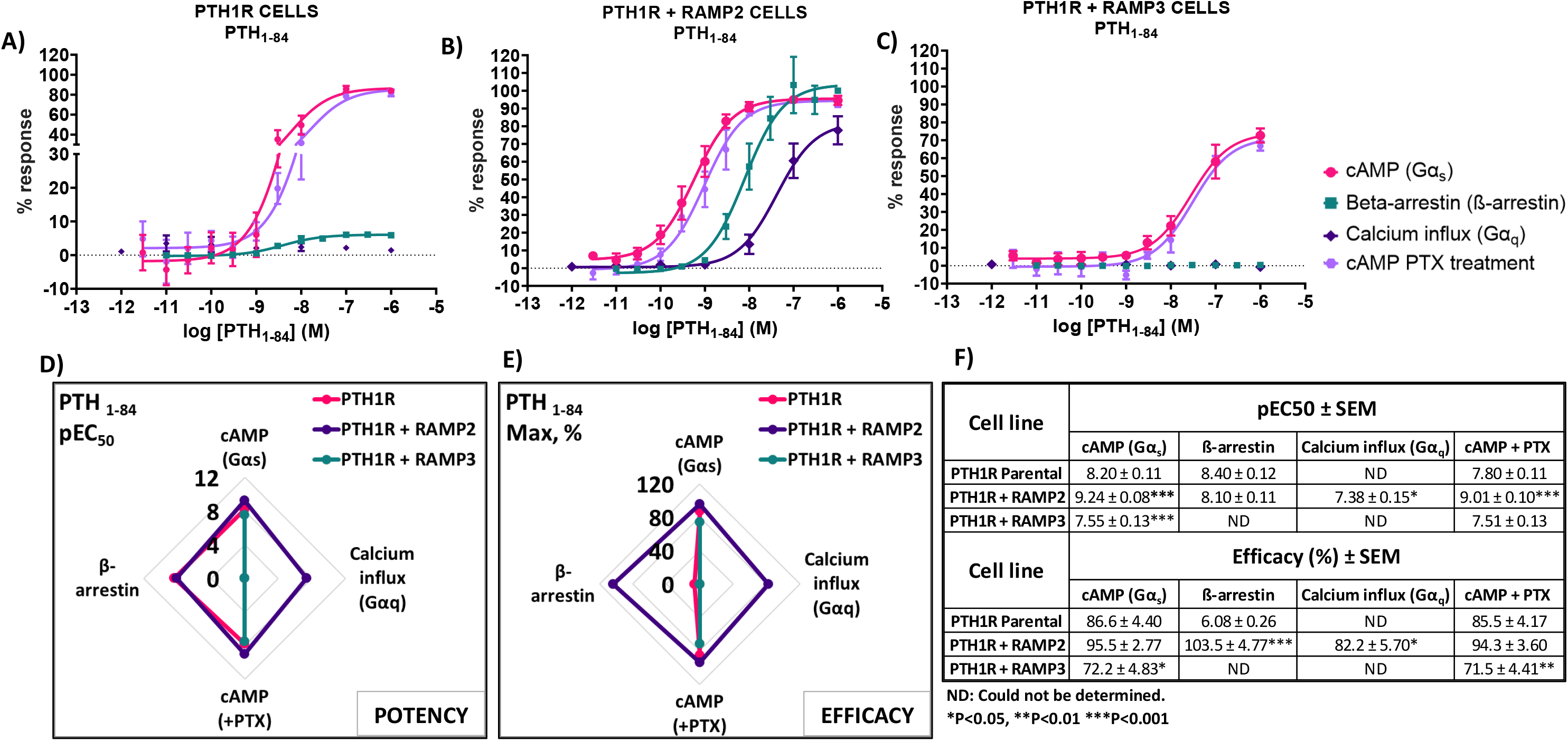
Consequences of RAMP2 and RAMP3 in the potency and efficacy of PTH (1-84) in different functional assays. Dose response curves of PTH (1-84) in different second messenger pathways, in cells overexpressing **a)** PTH1R alone, **b)** PTH1R with RAMP2 and **c)** PTH1R with RAMP3. Spider diagrams of the **d)** the potency and **e)** the efficacy of PTH (1-84) in those assays as extracted from the dose response curves. **f)** Table summarising the potency and the efficacy of PTH (1-84) and the statistical differences between the cells. Data are derived from curves constructed from at least 3-4 independently replicated studies, each consisting of 2 replicate measurements at each of 11 ligand concentrations. Data were analysed using comparison of fits (Graph Pad Prism) for non-linear regression curves, three-parameter logistic curve (*P<0.05, **P<0.01 ***P<0.001). All curves were expressed as a % of the positive controls/maximal response. Controls: cAMP studies forskolin (100μM), calcium influx studies ATP (100μM), β-arrestin-2 recruitment studies maximal response at highest dose (1μM).

### PTH (1-17)

Very similar effects were shown when we used a shorter chemically modified cyclic analogue of PTH (1-34), PTH (1-17) also known as ZP2307 (24). PTH1R RAMP2 cells showed a significant increase in both potency and efficacy in cAMP accumulation (Ga_s_) and Ga_i_, while PTH1R RAMP3 cells showed a significant decrease in both in cAMP accumulation (Ga_s_) and in only the efficacy of ZP2307 in Ga_i_ (Figure 9). In β-arrestin, like all the other peptides, ZP2307 had a significantly increased efficacy in PTH1R RAMP2 cells when compared to PTH1R alone, while PTH1R RAMP3 showed no response (Figure 9). Lastly, similarly to PTH (1-34), significant increase in efficacy but not potency was observed in calcium influx when PTH1R RAMP2 cells were stimulated with ZP2307, compared to PTH1R alone. PTH1R RAMP3 cells showed no response (Figure 9).

**Figure 9:**
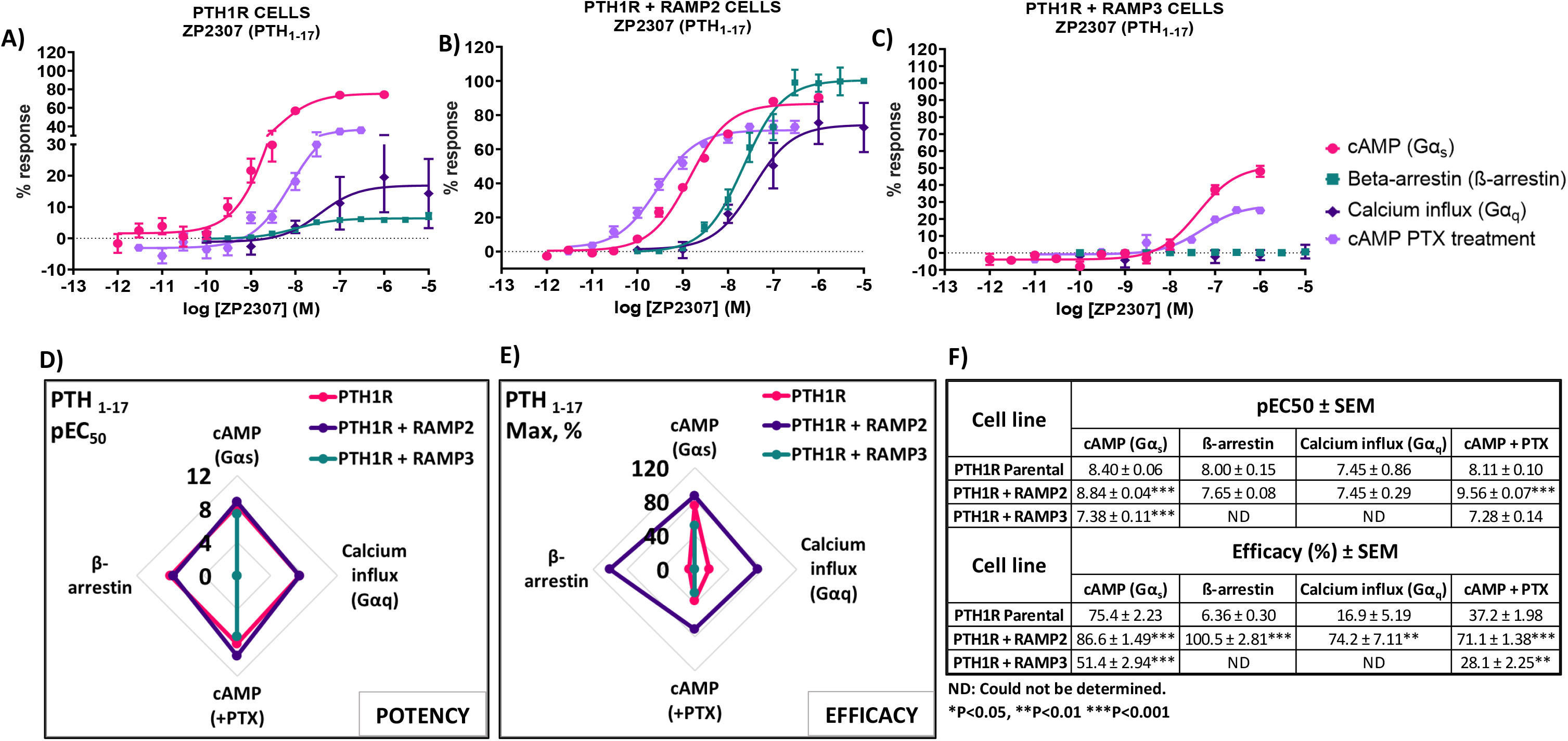
Consequences of RAMP2 and RAMP3 in the potency and efficacy of PTH (1-17)/ZP2307 in different functional assays. Dose response curves of PTH (1-17) in different second messenger pathways, in cells overexpressing **a)** PTH1R alone, **b)** PTH1R with RAMP2 and **c)** PTH1R with RAMP3. Spider diagrams of the **d)** the potency and **e)** the efficacy of PTH (1-17) in those assays as extracted from the dose response curves. **f)** Table summarising the potency and the efficacy of PTH (1-17) and the statistical differences between the cells. Data are derived from curves constructed from at least 3-4 independently replicated studies, each consisting of 2 replicate measurements at each of 11 ligand concentrations. Data were analysed using comparison of fits (Graph Pad Prism) for non-linear regression curves, three-parameter logistic curve (*P<0.05, **P<0.01 ***P<0.001). All curves were expressed as a % of the positive controls/maximal response. Controls: cAMP studies forskolin (100μM), calcium influx studies ATP (100μM), β-arrestin-2 recruitment studies maximal response at highest dose (1μM).

### PTHrP (1-108) and PTHrP(1-141)

We also tested effects of larger PTHrP analogues including the PTHrP (1-108) and full length PTHrP (1-141). Compared to the more widely studied PTHrP (1-34), these have statistically significant decreased potency and efficacy in activating PTH1R alone in cAMP accumulation studies (Figure 10 (d)). The presence of RAMP2 significantly increased the potency and efficacy of PTHrP (1-108) and the efficacy but not the potency of PTHrP (1-141) compared their activity on the PTH1R alone. However, here we show a statistically significant reduction in both potency and efficacy of cAMP activation in the presence of RAMP3 compared to stimulation of the PTH1R alone by PTHrP (1-108 & 1-141). Due to the limited availability of these peptides, we were unable to study the effects on other signalling pathways (calcium, βarrestin).

**Figure 10:**
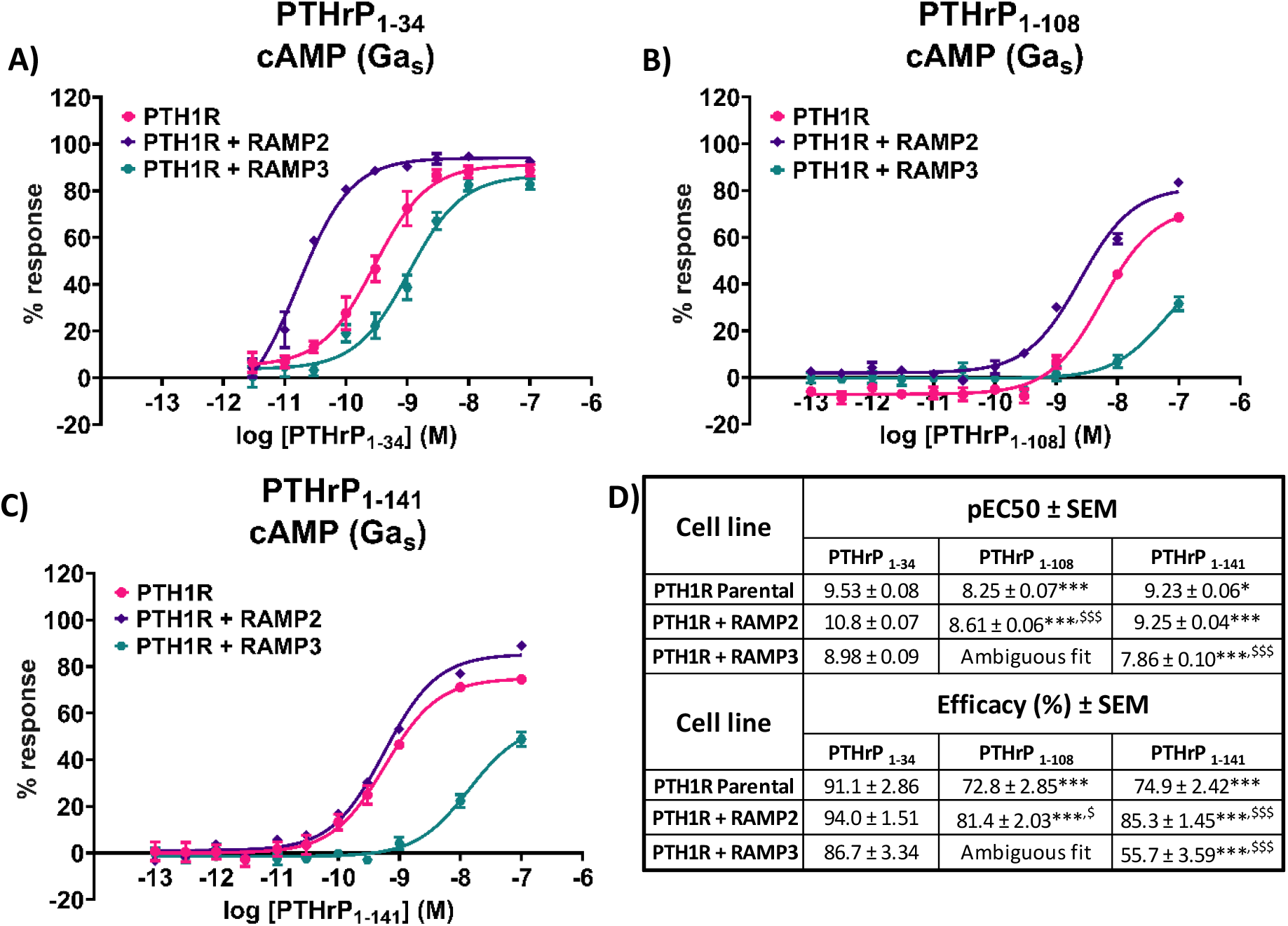
Effects of larger PTHrP analogues including the PTHrP (1-108) and the full length PTHrP (1-141) in cAMP accumulation studies compared to the more widely studied PTHrP (1-34). Dose response curves of **a)** PTHrP (1-34), b) PTHrP (1-108) and c)PTHrP (1-141) in cAMP accumulation studies, in cells overexpressing PTH1R alone, PTH1R with RAMP2 and PTH1R with RAMP3**. d)** Table summarising the potency and the efficacy of the different ligands and the statistical differences between them. Data are derived from curves constructed from at least 3-4 independently replicated studies, each consisting of 2 replicate measurements at each of 11 ligand concentrations. Data were analysed using comparison of fits (Graph Pad Prism) for non-linear regression curves, three-parameter logistic curve. All curves were expressed as a % of the positive control (forskolin (100μM)). *P<0.05, **P<0.01 ***P<0.001, for comparisons against PTHrP (1-34), ^$^P<0.05, ^$$^P<0.01 ^$$$^P<0.001, for comparisons against PTH1R alone cells.

## Discussion

In this study, we investigated the effects of receptor activity-modifying proteins (RAMPs) on the pharmacology of the parathyroid hormone 1 receptor (PTH1R) using a variety of PTH/PTHrP-related ligands. Our results demonstrate that RAMP2 and RAMP3 differentially interact with PTH1R and modulate its responses to these ligands. RAMP2 enhances β-arrestin recruitment and calcium signalling, while RAMP3 inhibits these pathways and appears to retain PTH1R intracellularly. We also show that the presence of RAMP2 differentially modulates the potency and efficacy of PTH/PTHrP related peptides in activating G proteins and recruiting β-arrestins. Furthermore, we report the effects of full-length PTHrP ligands, PTHrP (1-108) and PTHrP (1-141), on PTH1R signalling and the influence of RAMPs on these responses. Our findings highlight the complex role of RAMPs in modulating PTH1R function and suggest that targeting the PTH1R/RAMP2 maybe of potential therapeutic value.

Biased agonism is well-studied on PTH1R, with differing activation and duration in cAMP, calcium, and β-arrestins resulting in distinct physiological outcomes (25). The goal of translating biased agonist ligands into viable therapeutics is to increase bone mass while reducing calciotropic effects in conditions such as osteoporosis. However, less well studied is how accessory proteins, such as the RAMP family, play a role in biased agonism of PTH1R. PTH1R and RAMP interactions have previously been identified (10, 20). Nemec et al. undertook the first comprehensive pharmacological study of PTH1R and RAMP2 using PTH (1-34) and PTHrP (1-34) (11). In the present study, we investigated and expanded upon the possible effects of RAMPs on PTH1R pharmacology with a larger repertoire of PTH-derived ligands.

Our data show that both RAMP2 and RAMP3 form an interaction with PTH1R, but not RAMP1, consistent with observations by Harris et al. (20). Our FRET data confirm the preference of PTH1R to interact with RAMP2 and less so with RAMP3, while no interaction was shown with RAMP1 (Figure 2), as recently reported by Nemec et al. (11). We also observed different stoichiometric ratios of RAMP2 and RAMP3 interactions with PTH1R (Figure 2), similar to changes in GPCR/RAMP stoichiometry previously observed in the class C GPCR calcium-sensing receptor (26). The rationale for a FRET-based stoichiometric approach, as opposed to a pure FRET efficiency method, is that the latter is expressed in arbitrary units and cannot determine whether a low FRET signal is due to the absence of interaction between the components or to a local excess of donor and acceptor molecules. Using FRET stoichiometry, we can estimate the fraction of acceptor molecules in complex with donor molecules and the fraction of donor molecules in complex by measuring the donor fluorescence lost due to energy transfer (22). This eliminates the need for acceptor photobleaching to determine total donor concentrations and allows for repeated measurements from the same cells (22).

RAMP2 interaction with PTH1R does not alter cell surface expression of PTH1R, confirming observations by Christopoulos et al. (9). However, RAMP3 appears to almost completely ablate cell surface expression of PTH1R (Figure 3), possibly implying a retention preventing forward trafficking of PTH1R to the cell surface or an increase in PTH1R internalization/recycling, as recently shown by Mackie et al. (18). To confirm these further experimental studies, such as real time microscopy. It has previously been reported that PTH1R to localise in sub-cellular compartments including the nucleus (27). In recent years, it’s been increasingly recognized that GPCR signalling can continue after endocytosis. This phenomenon, called endosomal signalling, challenges the traditional view that GPCR signalling is mainly confined to the cell surface. The effects of the subcellular localization of PTH1R have previously been shown to affect ligand binding (28, 29). Studies have shown that PTHrP(1–36) or PTHrP (1–34) analog (Abaloparatide) primarily exerted their effects on the cell surface, whereas PTH(1–34) was more prone to endosomal internalization, resulting in an extended elevation of cyclic AMP levels in cells overexpressing PTH1R (29, 30).

Using antibody-capture SPA in COS-7, we showed that PTH1R with RAMP2 induces different responses to PTH (1–34) and PTHrP (1-34) compared to PTH1R alone (Figure 4), illustrating an example of ligand-induced functional selectivity responses for a receptor/RAMP complex, as observed previously (12, 31). RAMP2 increased PTH-stimulated maximal activation of Gα_s_ and Gα_i_ without changing Gα_q_. PTHrP induced a different pattern of changes, with PTHrP-stimulated Gα_s_ efficacy increased significantly without changes in Gα_i_, Gα_q_, or potency. PTH induced greater maximal activation of Gα_s_ and Gα_i_ than PTHrP and slight Gα_q_ activation that was absent with PTHrP. As PTH and PTHrP are known to bind to the same PTH1R but induce different tissue/organism effects (32, 33), this provides quantitative confirmation of ligand-induced functional selectivity at the level of G-protein activation by different physiological ligands binding to the same receptor (34). COS-7 cell model was used aspect of the study due to ability to express high levels of receptor at the cell surface and express a full complement of G-protein in the cell (31). COS-7 may not be the optimal cell system to perform studies on the PTH1R as receptor recycling and desensitisation (35) have not been observed in these cells, however, in this experiment we studied the activation of the G-proteins in isolated membrane fragments where internalisation and desensitisation is not studied. To address these issues and the whole functional assay CHO-K1 cells were used, which have previously been used to study internalization and desensitisation (36, 37).

Nemec et al. showed that RAMP2 specifically and selectively enhanced the activation kinetics of Gαs and Gαi3 proteins by PTH (1–34) (11), consistent with our data (Figure 4) showing larger maximal responses by PTH (1–34) on PTH1R Gα_s_ activation and Gα_i_ sensitivity in the presence of RAMP2. We also show a significant reduction in potency with no change in efficacy of Gα_q_ activation by RAMP2 (Figure 4), whereas others did not observe any differences. For PTHrP (1–34), we observed an increase in efficacy of only Gα_s_ with no alterations in Gα_i_ signalling (Figure 4), in contrast to observations where RAMP2 did not alter PTHrP (1-34) G protein activation by PTH1R. These discrepancies could reflect the different detection methods used.

The kinetics of cAMP activation play a crucial role in the differential effects of PTH and PTHrP. PTH (1-34) has been shown to induce prolonged cAMP activation compared to PTHrP (1-34) (29, 38). This prolonged cAMP activation is associated with the bone anabolic effects of PTH (1-34). In the presence of RAMP2, our results show that PTH (1-34) has higher potency and efficacy than PTH1R alone, suggesting an even longer duration/magnitude of cAMP activation. However, PTHrP (1-34) exhibits increased potency but unchanged efficacy in the presence of RAMP2, indicating a different pattern of cAMP activation kinetics possible be as a result of altered receptor internalisation and endosomal signalling (29). A limitation of our study is that we did not directly examine the kinetics of cAMP activation, which could be addressed in future studies to further elucidate the role of RAMPs in modulating the temporal aspects of PTH1R signalling.

The presence of RAMP2 significantly enhances the maximal response of β-arrestin recruitment by PTH1R for all ligands tested (Figures 6-9), consistent with the previous work by Nemec et al (11). Our data suggest that RAMP2 has a universal effect on augmenting βarrestin recruitment and PTH1R desensitization. Increased β-arrestin activity has been correlated with bone anabolic effects (8, 39), and the β-arrestin selective agonist DTrp12,Tyr34-bPTH (7–34) has been shown to increase bone formation without activating G proteins, inducing hypercalcemia, or increasing markers of bone resorption (40). These findings highlight the potential role of β-arrestin signalling in the regulation of bone metabolism and suggest that targeting β-arrestin recruitment through the PTH1R/RAMP2 complex may be a promising strategy for the development of novel bone anabolic therapies.

Our results also demonstrate that PTH (1–34) induces changes in the efficacy and/or potency of all downstream second messenger used (cAMP, calcium and β-arrestin), while PTHrP (1-34) elicits a dramatic increase in β-arrestin recruitment but no change in other second messengers. Interestingly, PTHrP (1-34) has been shown to have comparable bone anabolic effects compared to PTH (1-34) (41, 42). Our data may suggest that the bone anabolic effects of PTH (1–34) and PTHrP (1-34) could be mediated through the PTH1R/RAMP2 complex rather than PTH1R alone, with similar effects observed for PTH (1-84). Additionally, the differential effects of PTH (1-34) and PTHrP (1-34) on G protein and β-arrestin signalling in the presence of RAMP2 could provide new insights into the mechanisms underlying their distinct bone anabolic properties. Our findings suggest that the PTH1R/RAMP2 complex may be a key mediator of the bone anabolic effects of PTHrP (1-34) and potentially other PTH1R ligands. Further research is needed to elucidate the precise role of RAMP2 in regulating PTH1R signalling and its implications for bone physiology and disease.

The effects of RAMP3 on PTH1R have not been reported previously. As described above, we see an interaction between PTH1R and RAMP3. The cell-surface ELISA data shows reduced PTH1R expression, which may suggest a receptor retention effect by RAMP3, yet we still see responses by PTH1R/RAMP3 albeit, reduced, compared to PTH1R/RAMP2. β-arrestin and calcium signalling are absent in the presence of RAMP3 compared to PTH1R/RAMP2 and PTH1R alone. In the presence of RAMP3, the lack of cell surface trafficking and second messenger signalling is inconsistent and may be due to the difference in cellular backgrounds used in this study. However, there have been reports of intracellular PTH1R activation (43). Nonetheless, RAMP3 has also been reported to be an early response gene to PTH stimulation, further suggesting a potential important regulatory function of the PTH1R/RAMP3 interaction (44).

During this study we were also able to test less commonly explored PTHrP (1–108) (full-length analogue) and PTHrP (1-141) (full-length) peptides (45). These full-length PTHrP ligands have not been widely studied due to challenges in their synthesis. Compared to the more widely studied PTHrP (1–34), these have lower potency and efficacy in activating PTH1R in cAMP accumulation studies. The presence of RAMP2 increased their potency and/or efficacy when compared to PTH1R alone. However, consistent with our previous observations, RAMP3 reduces the response of PTH1R to these ligands when compared to PTH1R alone. Due to the limited availability of these peptides, we were unable to study the effects on other signalling pathways (calcium, β-arrestin). This strengthens the implication that RAMP2 plays a broad role in modulating PTH1R pharmacology across a variety of ligands.

The study by Kadmiel et al. provides valuable *in-vivo* evidence supporting the physiological relevance of RAMP2-GPCR interactions beyond the canonical AM-CLR signalling paradigm (21). The reduced PTH1R expression in Ramp2-/placentas and the blunted response to very large doses (500μg/kg) of systemic PTH administration in Ramp2+/adult females complement our *in-vitro* data demonstrating that RAMP2 modulates PTH1R signalling (21). These data suggest how these *in-vitro* changes in ligand bias reported here may influence *invivo* functions, however, this needs to be explored in more detail using physiological relevant levels of PTH and/or PTHrP.

In summary, our findings highlight the complex role of RAMPs in modulating PTH1R signalling and function. We show that RAMP2 and RAMP3 differentially interact with PTH1R and modulate its responses to a diverse range of PTH/PTHrP-related ligands. The presence of RAMP2 enhances β-arrestin recruitment and calcium signalling, while RAMP3 appears to reduce cell surface expression of PTH1R, and subsequently reduced PTH1R signalling is observed. The differential effects of PTH (1–34) and PTHrP (1-34) on G protein and β-arrestin signalling in the presence of RAMP2 could provide new insights into the mechanisms underlying their distinct bone anabolic properties. Moreover, our data also warrants detailed understanding of whether the kinetics of cAMP activation is, differentially modulated by RAMP2 for PTH (1-34) and PTHrP (1-34) and can likely contribute to their divergent physiological effects. One constraint in our research is understanding if the binding affinities of these ligands are altered via allosteric modulation by RAMPs, especially RAMP2, and whether it may impact the interpretation of the signalling data. Overall, our data suggest that targeting the PTH1R/RAMP2 complex may be a promising strategy for the development of novel bone anabolic therapies by potentially leveraging functional selectivity. Further research using functional readouts in primary cells, and appropriate animal models, including knockout mice, will be crucial to elucidate the physiological relevance of these findings and their potential therapeutic implications.

## Materials and methods

### Materials

Reagents were purchased from respective manufacturers: Ham’s F12-K (Kaighn’s) Medium; RPMI Medium; sodium pyruvate; penicillin/streptomycin; foetal bovine serum (FBS); OptiMEM™ (Reduced Serum Medium); Lipofectamine 3000 (GIBCO-Invitrogen-Life Technologies, Carlsbad, CA); AssayComplete™ Cell Culture Kit-107 (DiscoverX, California, US). ATP, forskolin, IBMX and Pertussis toxin (Sigma Aldrich, St. Louis, MO); G418 (Fisher Scientific, Loughborough, UK); rabbit anti-goat-HRP antibody and goat anti-mouse-HRP antibody (Dako, Denmark). LANCE cAMP Detection kit (Perkin Elmer, Massachusetts, US); FLIPR Calcium 6 Evaluation Kit (Molecular Probes, Oregon, US); PathHunter® Detection Kit (DiscoverX, California, US). CHO-K1 and COS-7 cell lines (ATCC, Virginia, US); PathHunter CHO-K1 PTH1R β-arrestin cell line (DiscoverX, California, US). PTH (1-34); PTH (1-84); PTHrP (1-34) (Bachem Holding, Bubendorf, Switzerland). PTH (1-17) also known as ZP2307 was a gift from Dr Rasmus Just of Zealand Pharma. Truncated PTHrP peptides were those developed and produced by Professor Jack Martin (45).

### Cell transfections and Cell line generation

COS-7 cells were grown to confluency in DMEM with GlutaMAX^TM^, supplemented with 10% FCS and 1x penicillin/streptomycin, in a 5% CO2 incubator at 37°C. The cells were harvested using trypsin/EDTA (Sigma), washed with PBS, and resuspended in electroporation buffer (composition [mM] 20 HEPES, 135 KCl, 2 MgCl_2_, 2 ATP, 5 glutathione, 0.5% Ficoll 400 adjusted to pH 7.6 using KOH) at a concentration of ∼4 million cells into 4mm gap electroporation cuvettes (York Biosciences, UK) and the required DNA was added (5μg PTH1R, 15μg RAMP constructs). The cells were then electroporated at 0.25 kV and 960 µF using a Gene Pulser (Biorad) and cultured for 48hr.

CHO-K1 PTH1R β-arrestin cells were grown in AssayComplete™ Cell Culture Kit-107, containing all necessary supplements and antibiotics, at 37°C in 5% CO_2_ and were subcultured in a 1:10 ratio every 3-4 days. To be used in the functional assays (cAMP accumulation, β-arrestin recruitment and calcium mobilisation) cells were transfected with Ctagged Ceruleaun RAMPs (RAMP1-3) constructs using Lipofectamine 3000 following the manufacturer’s guidelines. Cells were selected using 0.5 mg/mL G418 48 hours after transfection and cultured in the aforementioned growth media for 1 week. RAMP expressing cells were validated using fluorescence imaging using the EVOS microscope and population enrichment by fluorescence-assisted cell sorting using the FACS Aria II (BD Biosciences, New Jersey, US). This was done twice (two separate sorts).

### Membrane preparations

Cell membrane extractions were performed using COS-7 cells transfected with PTH1R and different RAMPs (described above) and used in Scintillation proximity assay. At 48hours post transfection the cells were homogenized using a Dounce homogenizer using ice cold PBS and centrifuged at 300g for 10 minutes at 4°C in a final volume of 40ml. The supernatant was collected in a fresh tube and centrifuged at 50,000g for 25 minutes at 4°C. The resulting pellet was resuspended in ice cold SPA buffer (50mM HEPES, 100mM NaCl, 5mM MgCl2, 0.5% BSA, pH 7.4). Total protein concentrations were measured using Bicinchoninic acid assay (Sigma).

### Preparation of constructs for FRET and COS-7 cell transfection

FRET studies were performed with minor modifications to previously published methods (26). Citrine or Cerulean cDNAs were engineered into a pcDNA3.1 vector (Invitrogen) between the Not1 and Xho1 restriction enzyme sites. RAMPs and PTH1R were engineered into pcDNA3.1 Cerulean and Citrine vectors respectively excluding their stop codons, between the Kpn1 and Not1, and HindIII and Not1 restriction enzyme sites, so that the fluorophores were present at the C-terminal of RAMP/PTH1R. As a negative control, pcDNA 3.1 containing Citrine alone were co-transfected with a pcDNA3.1 RAMP Cerulean vector. As a positive control, we created a pcDNA3.1 vector containing a Cerulean cDNA fusion construct followed by 18 amino acid linker sequence and then Citrine cDNA.

COS-7 cells were grown to confluency and harvested using trypsin/EDTA (Sigma), washed with PBS, and resuspended in electroporation buffer (composition [mM] 20 HEPES, 135 KCl, 2 MgCl2, 2 ATP, 5 glutathione, 0.5% Ficoll 400 adjusted to pH 7.6 using KOH) at a concentration of ∼3.5– 4 million cells into 4 mm gap electroporation cuvettes (York Biosciences, UK) and the required concentration of DNA was added (10 µg receptor, 15 µg RAMP constructs). The cells were then electroporated at 0.25 kV and 960 µF using a Gene Pulser (Biorad) and cultured for 72 hr in 35 mm glass-bottom plated (Ibidi, München) after which they were fixed with 4% PFA and mounted with Mowiol. COS-7 cells were transfected with C-tagged Citrine PTH1R and Ctagged Ceruleaun RAMP in pcDNA 3.1 vector and grown in 35 mm glass-bottom plates (Ibidi, München) then fixed and mounted. As a positive control, fusion of Cerulean-Citrine was created in pcDNA3.1 vector. Cells were excited and images were captured for analysis.

Images were captured using a Zeiss Plan apo 63×/1.4 oil immersion lens on a Zeiss LSM 510 inverted laser scanning confocal fluorescence microscope fitted with an argon laser at room temperature. Confocal images of the fluorescent proteins were acquired using an argon laser together with an HFT458/514 nm dichroic, a NFT515 nm beam splitter, pin hole set to 496µm, detector gain 550 and individually as a separate channel under the following conditions: Cerulean was excited using the 458 nm laser line with a 100% laser intensity and a band pass BP480–520 emission filter; Citrine was excited using the 514 nm laser line attenuated to 20% laser intensity and a band pass BP535–590 emission filter; FRET was excited using the 458 nm laser line with a 100% laser intensity and a BP480–520 emission filter. All fluorescence channels were scanned and collected, line by line with a mean of 1.

Cerulean and Citrine fluorescence bleed-through into the FRET channel were calculated using FRET and co-localization analyser plugin for ImageJ (46). NFRET calculations for FRET efficiency for sensitized emission were done using pixel-by-pixel analysis by PixFRET plugin for ImageJ (47). The threshold for pixel intensity to be included in analysis was set to 1.5 times the background intensity. The following equation was used to calculate FRET efficiency:

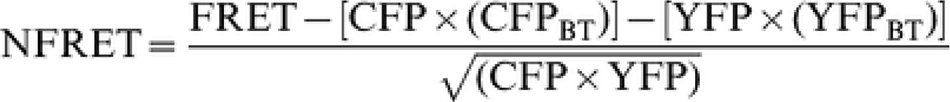

### BT= bleed through

The correction factors calculated were: β (proportionality constant relating donor fluorescence detected at the acceptor emission relative to that detected at the donor emission): 0.31, α (proportionality constant relating acceptor fluorescence at the acceptor excitation to the donor excitation): 0.126, γ (ratio of the extinction coefficient of the acceptor to the donor at the donor excitation): 0.3, ξ (proportionality constant relating the sensitized acceptor emission to the decrease in donor fluorescence due to FRET): 0.2.

Cell surface FRET was separated from whole cell FRET by constructing a series of 50-pixel diameter dots around the cell surface of the raw acceptor image using the selection tool of Image J. Each dot was taken as a ROI and the combined ROIs for each image were used to calculate mean membrane NFRET and stoichiometry values. All FRET-based stoichiometric analysis was performed as previously described (22) using ImageJ software.

### Cell surface expression

Cells (CHO-K1 parental or stable cells generated as described in Cell Line Generation section above) were seeded at 150,000 cells/well into 24-well plates coated with poly-D-lysine. Following 48 hours growth in complete growth media, media was replaced with 4% formaldehyde for 15 min to fix the cells. Cells were washed 3 times with 500 µL phosphatebuffered saline (PBS) and incubated with 1% BSA in PBS for 45 min to prevent nonspecific antibody binding. To determine receptor expression, 250 µL of primary antibody (mouse anti-PTH1R, (ab104832 (1mg/ml), Abcam, Cambridge, UK)), at 1:3000 concentration in 1% BSA PBS was added for 1 hour. Cells were washed 3 times with 500 µL PBS before the addition of the secondary antibody (HRP-conjugated anti-mouse IgG (1.5mg/mL) (Dako, Denmark)) diluted 1:2000 in 1% BSA PBS for 1 hour. Following 3 further washes with PBS, HRP activity was determined using TMB Substrate Solution (Thermo Scientific, Massachusetts, US) according to the manufacturer’s instructions.

### Scintillation proximity assay for G-protein activation

Receptor/G-protein activation profiles were determined using a scintillation proximity assay as described previously (48). Briefly, COS-7 cells were transfected with different native untagged human receptors alone or in combination with native untagged human RAMPs1-3 in pcDNA3.1. Receptor concentrations were determined by radioligand binding studies and Western blotting (not shown). For the SPA assay, membranes were incubated with different concentrations of ligands with [35^S^] GTPγ^S^. Polyclonal antibodies to different G-proteins were added (Santa Cruz). Scintillation proximity assay beads coated with secondary antibody were added and measured in a scintillation counter. Dose response curves were generated from no fewer than 2 independent replicates at 8 different ligand concentrations, in 3 independent studies for each receptor and RAMP comparison.

### Intracellular calcium mobilisation assay

To assess the consequences of RAMP expression in calcium influx, calcium mobilisation assays were performed in 96-well black, clear-bottom plates (Corning, USA). 48 hours before the assay was performed, CHO-K1 PTH1R β-arrestin cells expressing the different RAMPs (generated as explained above) were seeded at 10,000 cells/well in standard growth medium to give 80% confluency at time of assay. The medium was replaced with 1% FBS medium 24 hours prior to stimulation. After thawing and equilibrating the 10x Calcium 6 assay reagent to RT, it was dissolved in 10 mL (1:10) of loading buffer (1x HBSS buffer, 20 mM HEPES, 10 mM CaCl2, and pH adjusted to 7.4). Probenecid was added to loading dye to give final in-well concentration of 2.5 mM (this prevents the release of the dye from the cells back into to the medium). 100 µL of 1x Calcium 6 loading dye was added to all wells and incubated for 2 hours at 37oC, 5% CO2. All peptide ligands used were diluted in 1x loading buffer. Following incubation, the plate was then transferred directly to the Flexstation3 assay plate reader (Molecular Devices, California, US) and was allowed to equilibrate at 37°C for 10 mins. Traces were collected for 300 seconds, including a 50 second baseline read prior to peptide addition. All intra-experimental traces were collected in duplicate. The fluorescence values after exposure were subtracted by the basal fluorescence value before exposure and the data were normalised using ATP (1μΜ) stimulated controls as 100% response. Dose response curves were analysed using three-parameter logistic curve to determine EC_50_ values (Graphpad Prism 9 and 10).

### Time-resolved fluorescence resonance energy transfer (TR-FRET) cAMP accumulation

To assess the consequences of RAMP expression in cAMP accumulation, the total cAMP was measured using the TR-FRET LANCE cAMP detection kit (PerkinElmer, AD0264), according to the manufacturer’s directions. The assay was performed using CHO-K1 PTH1R β-arrestin cells expressing the different RAMPs (generated as explained above). Aliquots of frozen cells (2 × 10^6^ each) were thawed and prepared in warm stimulation buffer (1 × HBSS, 5 mM HEPES, 0.5 mM IBMX, and 0.1% BSA). Alexa Fluor antibody (1:100 concentration) was added to the cell suspension and cells were plated (2,500 cells, 6 μL) in a 384-well white opaque microtiter plate (OptiPlates, Perkin Elmer, 6007299). Cells were incubated with serial dilutions (3 μL) of the peptide ligands (agonist) for 30 minutes at RT. Subsequently, 12 µL detection mix (Europium-Chelate streptavidin/biotinylated cAMP) was added to stop the reaction and induce cell lysis. TR-FRET was detected after an hour incubation by an EnSight multimode Plate reader (Perkin Elmer), at; 320/340nm excitation and 615/665nm emission. Data were normalised to a forskolin (100μM) only control as 100% cAMP accumulation. Dose response curves were analysed using three-parameter logistic curve to determine EC_50_ values (Graphpad Prism 9 and 10).

### Pertussis toxin treatment

For investigation of Gα_i_ modulation cells were pre-treated with growth media supplemented with 200 ng/µL pertussis toxin (PTX) (Sigma-Aldrich, USA). Following an overnight incubation with PTX, cells were frozen down and were used following the cAMP detection procedure above.

### Beta (β)-arrestin recruitment

To assess the consequences of RAMP expression in the recruitment of β-arrestin (β-arrestin2 isoform) the PathHunter® Detection Kit was used. The assay was performed following the manufacturer’s instruction and by using CHO-K1 PTH1R β-arrestin cells expressing the different RAMPs (generated as explained above). More specifically, 20 μL (5,000 cells/well) was added in a 384-well white opaque microtiter plate. Cells were then incubated with serial dilutions (5 μL) of the peptide ligands (agonist), prepared in 1x HBSS + 20mM HEPES buffer, for 90 minutes at RT. 12.5 μL Working Detection Solution (mix 19 parts of cell assay buffer, 5 parts of Substrate Reagent, and 1 part of Substrate Reagent 2) was then added to the wells. Chemiluminescence was detected an hour after using EnSight multimode Plate reader. Data were normalised to the maximal response at highest ligand dose (1μM) as 100% and to no ligand as 0% response. Dose response curves were analysed using three-parameter logistic curve to determine EC_50_ values (Graphpad Prism 9 and 10).

## Acknowledgements

We would like to thank the following for their assistance with these studies: BBSRC (Industrial Partnership Award, 117120) for funding and supporting these studies; Dr Rasmus Just of Zealand Pharma for his generous gift of PTH (1-17) also known as ZP2307; and Professor Jack Martin from the University of Melbourne for his generous gift of the PTHrP peptides (PTHrP (1-108) and (1-141)) and critical review of this manuscript.

## Authors contributions

T.M.S., and G.O.R., idea conception, P.A., A.J.D., D.J.R., and G.O.R. designed research; P.A., A.J.D., D.J.R., E.R.L., and G.W.S. performed research; P.A., A.J.D., A.B.A.J., D.J.R. and G.O.R. analysed data; P.A., A.J.D., A.B.A.J., T.M.S., and G.O.R. wrote the manuscript; P.A., A.J.D., A.B.A.J., D.J.R., E.R.L., G.W.S., T.M.S., and G.O.R. reviewed the manuscript.

